# Mitochondria redistribution organizes the immunosuppressive tumor ecosystem

**DOI:** 10.1101/2025.11.11.687895

**Authors:** Azusa Terasaki, Alexis T. Weiner, Yuhao Tan, Viktoria Szeifert, Keshav Bhatnagar, Cara C. Rada, Vishnu Shankar, Courtney Kernick, Mahnoor Mahmood, Lukas Wiggers, Viviana R. Rodrigues, Payam A. Gammage, Theodore Roth, Jeffrey D. Axelrod, Edgar Engleman, Bo Li, Derick Okwan-Duodu

## Abstract

Hostile conditions in the tumor microenvironment restrict cellular respiration, yet mitochondrial metabolism remains indispensable for tumor growth and the activity of immunosuppressive cells. How tumor ecosystems sustain mitochondrial output has been unclear. Here, we show that cancer cells resolve this paradox by acting as hubs of intercellular mitochondrial redistribution. Using mitochondrial reporter systems, we demonstrate that cancer cells import host-derived mitochondria, integrate them into their endogenous network, and subsequently relay these hybrid organelles to neighboring immune cells. Mitochondria redistribution reprograms recipient neutrophils, macrophages, and CD4+ T cells into highly suppressive states but drives CD8+ T cell exhaustion. Within cancer cells, fusion of incoming mitochondria induces filamentous P5CS assembly, enhances biosynthetic output, and enables the refurbishment of damaged organelles into fully functional units. Disrupting mitochondrial redistribution collapses the immunosuppressive ecosystem and impairs tumor growth. Thus, cancer cells do not hoard resources but orchestrate a redistribution program that fortifies their own metabolic resilience, derails anti-tumor immunity, and sustains immunosuppressive partners.

**HIGHLIGHTS:**

- Tumor cells regulate their ecosystem by redistributing mitochondria
- Redistributed mitochondria expand immunosuppressive cells but exhausts CD8+ T cells
- Mitochondria fusion within cancer cells, which precedes redistribution, optimizes metabolic output by triggering conformational changes in P5CS
- Mitochondria fusion allows cancer cells to incorporate and refurbish seemingly incompetent host-derived mitochondria, improving efficiency in the tumor ecosystem

## INTRODUCTION

The tumor microenvironment (TME) imposes severe constraints on oxidative metabolism^1^. Hypoxia, abnormal vasculature, metabolic competition, and the accumulation of multiple inhibitory metabolites collectively compromise mitochondrial respiration and are thought to underlie failure of anti-tumor immunity and immune-directed therapies^2–6^. Yet, paradoxically, tumor cells and immunoregulatory populations within the same hostile TME maintain robust mitochondrial activity, supporting their persistence and function^7–9^. For instance, despite the canonical Warburg effect^10^, the vast majority of metabolism in cancer cells is mitochondria-derived^8,11^. Moreover, regulatory T cells and macrophages rely on mitochondrial metabolism to execute most of their immunosuppressive functions^12,13^. Because solid tumors rarely regress spontaneously ^14^, these observations point to an underlying mechanism by which tumors maintain a mitochondrial metabolic ecosystem that selectively supports cancer cells and their immunosuppressive partners. Precisely how this is achieved remains unresolved.

Emerging studies indicate that cancer cells can supplement their metabolic needs by directly harvesting exogenous mitochondria from the host ^15–19^, a process that has been proposed to boost ATP production in cancer cells^15,20,21^. However, the fate of these exogenously acquired mitochondria and their broader consequences for the TME are unknown. Moreover, it remains unclear how a relatively small input of transferred exogenous mitochondria elicits dramatic bioenergetic output in recipient cells. Here, leveraging tractable mitochondrial reporter systems across murine models and patient samples, we unexpectedly found that cancer cells fuse acquired exogenous mitochondria with their own and ultimately redistribute them to fuel immunosuppressive cells. In the process, cancer cells themselves benefit by optimizing their metabolic capacity through architectural rearrangement of P5CS, a critical enzyme for macromolecular biosynthetic capabilities of cancer cells^22^. These findings uncover a previously unrecognized regulatory axis in which cancer cells act not merely as resource competitors, but as active organizers of a mitochondrial economy that fuels immune suppression and tumor progression.

## RESULTS

### Cancer cells redistribute mitochondria

To study intercellular mitochondrial transfer, we developed a flow cytometry and confocal microscopy platform in which cancer cells were co-cultured with leukocytes from PhaM^excised^ mice, which express a mitochondria-targeted dendra2 (mtD2) reporter^23^. In agreement with prior reports^16,18,24,25^, we confirmed that tumors (B16 melanoma or E0771 breast) rapidly acquired mtD2 protein during co-culture, indicating mitochondria transfer from immune cells (**Fig S1A** and **Fig S1B**). Three observations raised the possibility that cancer cells do not permanently retain exogenous mitochondria. First, after mitochondria transfer to cancer cells, the mtD2+ signal declined faster than could be explained by proliferative loss from cell division, consistent with elimination or redistribution (**Fig S1C**). Second, across tumor: leukocyte co-culture ratios (1:1 to 1:10), the fraction of mtD2+ tumor cells increased, but per-cell mtD2 intensity remained constant (**Fig. S1D**), suggesting a set-point for mitochondrial content. Third, live-cell imaging confirmed early mtD2 uptake by melanoma cells, followed by return of signal to immune cells (**Supplementary Video**).

To formally test whether cancer cells redistribute exogenously acquired mitochondria, we leveraged the CD45.1 and CD45.2 allelic variants mouse systems^26^, such that labeled mitochondria originating from CD45.2 cells (CD45.2^mtD2+^) can be unambiguously tracked in mitochondria-unlabeled CD45.1 cells (CD45.1^mtD2-^). CD45.1^mtD2-^:CD45.2 ^mtD2+^ co-cultures result in minimal mtD2 transfer to CD45.1 cells. In triple co-culture with B16 melanoma, CD45.1 cells became mtD2+ (**Fig 1A,B**). However, when B16 cancer cells are indirectly present (that is, seeded in Boyden chamber with 3 µm filters during the CD45.1^mtD2-^:CD45.2^mtD2+^ co-culture to prevent direct contact), there was minimal mitochondria transfer to CD45.1 cells (**Fig 1C**), suggesting that cancer cells act as direct conduits for mitochondria transfer between immune cells. To definitively demonstrate that cancer cells directly transfer mtD2+ exogenous mitochondria, we first co-cultured CD45.2^mtD2+^ leukocytes with B16 and sort-purified mtD2+ cancer cells (**Fig 1D**). In subsequent direct co-culture with CD45.1^mtD2-^, tumor cells transferred some of the previously acquired mtD2 to CD45.1 cells (**Fig 1E**). Confocal imaging confirmed cancer cell redistribution of mtD2+ mitochondria to CD45.1-derived neutrophils, macrophages, and T cells (**Fig 1F**).

**Figure 1.**
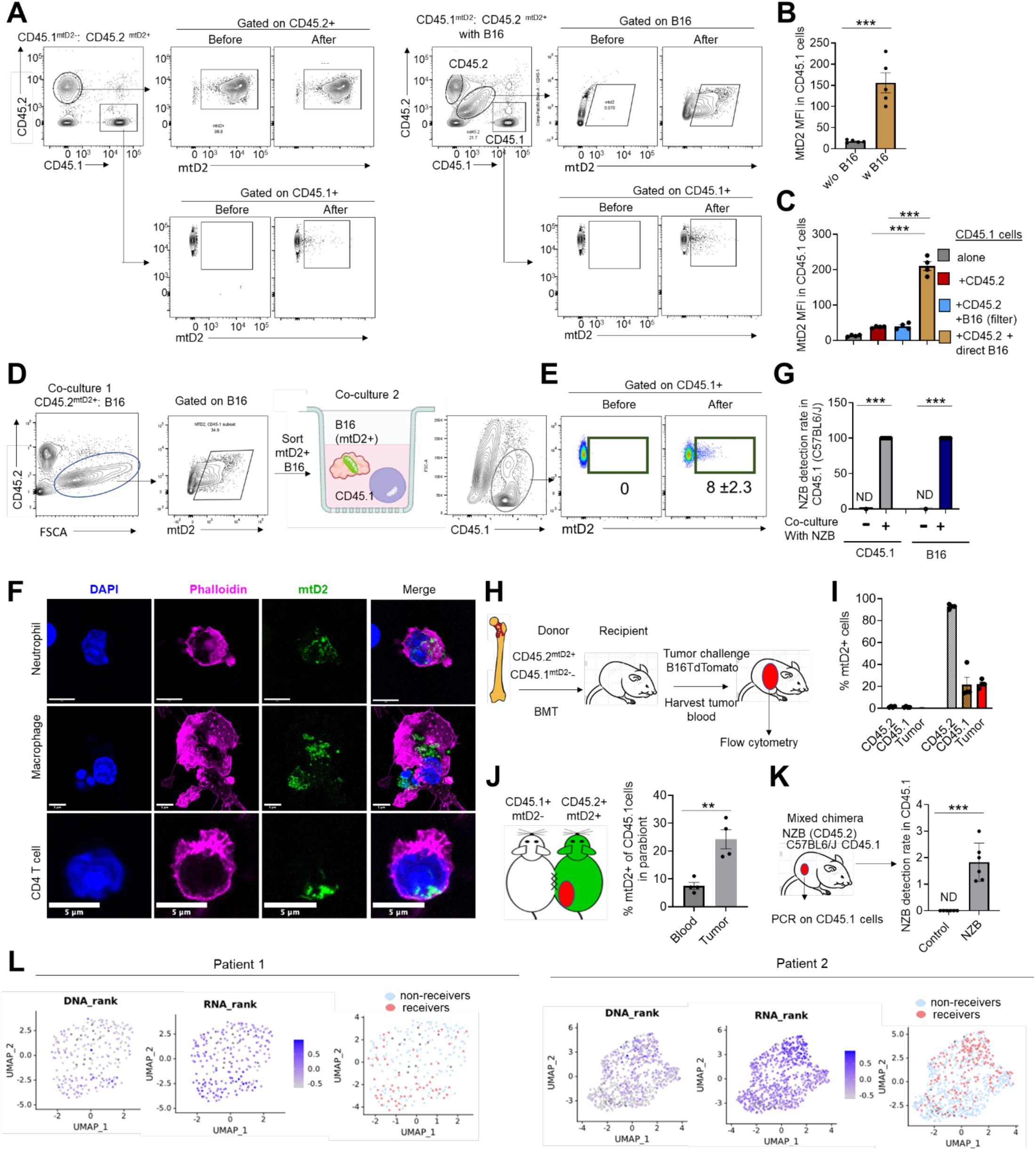
Cancer cells redistribute exogenously acquired mitochondria. **A**) Gating strategy for the detection of transferred CD45.2-derived, labeled mitochondria (mtD2) to cancer cells and to CD45.1 immune cells. **B**) mtD2 mean fluorescence intensity in CD45.1 cells cultured with CD45.2^mtD2+^ cells alone, or with CD45.2^mtD2+^ cells along with cancer cells (B16 melanoma). n=6 mtD2 mice and CD45.2 recipients, 3 independent times. **C**) Detection of mtD2 in CD45.1 cells cultured alone, or in the presence of CD45.2^mtD2+^ cells along with the direct or indirect cancer cell co-culture. **D**) Schematic for the culture of B16 melanoma with CD45.2^mtD2+^ cells, followed by sorting of mtD2+ cancer cells and subsequent co-culture with CD45.1 leukocytes. **E**) mtD2 detection in CD45.1 cells co-cultured with mtD2+ tumor cells. **F**) Representative immunofluorescence images of CD45.1 immune cells (neutrophils, macrophages and lymphocytes) that acquired mtD2 protein from cancer cells previously co-cultured with CD45.2^mtD2+^ cells. Scale bar= 5 µm. **G**) PCR detection rate of NZB mtDNA in CD45.1 cells that were co-cultured with cancer cells that had interacted with NZB leukocytes. **H**) Schema for the establishment of mixed chimeric mice from CD45.1^mtD2-^ and CD45.2 ^mtD2+^ donors. n=6 chimeras. **I**) Detection of mtD2 in CD45.1 cells in the mixed chimeric mice challenged with B16 tumors. **J**) Detection of mtD2 in CD45.1 cells from CD45.2^mtD2+^ and CD45.1^mtD2-^ parabionts challenged with tumor. n= 4 parabionts. **K)** PCR detection of NZB mtDNA heteroplasmy in CD45.1 cells in the tumor microenvironment of NZB/C57 mixed chimeric mice implanted with tumors. n= 6 mixed chimeras per group. **L)** Computational inference of redistributed mitochondria from CD8^+^ T cells to macrophages in the tumor microenvironment of human patients with basal cell carcinoma and melanoma from GSE123814. *p<0.05, **p<0.01, ***p<0.001 by unpaired Student’s t test except c (ANOVA).

We repeated these co-culture experiments using other cancer cell types, such as E0771 breast cancer. Moreover, this time we tracked mitochondria from CD45.2 (using CD.45.2^mtD2+^) to CD45.1 (CD45.1^mtD2-^) to demonstrate that the findings are allele variant-independent. The results were identical to what we described for B16: in the direct presence of E0771, mitochondria from CD45.2 can be redistributed to CD45.1 cells (**Fig S1E**).

For genetic confirmation of mitochondria redistribution by cancer cells independent of any potential dendra2 artifacts, we co-cultured B16 with leukocytes from NZB/BINJ mice, which are on a CD45.2 background. We subsequently co-cultured the tumor cells with CD45.1 cells (from C57BL6/J). As the NZB exhibits defined mitochondria polymorphisms compared to the C57BL6/J reference mtDNA genome^27^, ARMS-PCR can be performed on sorted CD45.1 cells which, if positive, indicates the acquisition of mtDNA from NZB^28^. In all cases, NZB mtDNA was detected in CD45.1 and B16 cells (**Fig 1G**). This approach further supported that cancer cells acquire exogenous mitochondria and redistribute them to cells in their microenvironment.

### Mitochondria redistribution *in vivo* by mice and human cancers

We next tested mitochondria redistribution by tumor cells *in vivo* using various complementary approaches. First, we established mixed chimeras, in which lethally irradiated C57BL6/J mice were reconstituted with hematopoietic cells from CD45.2^mtD2+^ and CD45.1^mtD2-^ donors (**Fig 1H**, and **Fig S1F**), enabling the tracking of dendra2 from CD45.2 to CD45.1 cells upon tumor implantation. These chimeric mice were then challenged with subcutaneous tumor implantation of B16Tdtomato. After tumors were established (∼day 14), they were excised and processed for flow cytometric analysis of live cells to examine the acquisition of mitochondria. As expected, tumor cells became positive for mtD2+, suggesting the horizontal transfer of mitochondria from CD45.2 immune cells. Notably, mtD2 was also detected in CD45.1 immune cells in the TME (**Fig 1I**). Second, we performed parabiosis of CD45.2^mtD2+^ and CD45.1^mtD2-^ mice. After chimerism was established, tumors were implanted and we confirmed the detection of mtD2+ mitochondria from CD45.2 cells to CD45.1 infiltrating the TME (**Fig 1J**). Third, we established a mixed chimera of CD45.2^NZB^ and CD45.1^C57BL6/J^. Animals were challenged with tumors. NZB mtDNA was detected in tumor cells and in the CD45.1 cells (**Fig 1K**), paralleling the *in vitro* results. Collectively, the results indicate that tumors can acquire mitochondria from the host and and disperse them to other immune cells.

Cell type–restricted mtD2 expression further demonstrated mitochondria redistribution between immune lineages. In PhAM^floxed^ mice crossed to CD8^Cre^, mtD2 was initially confined to CD8+ T cells (**Fig S1G**). In the presence of B16 tumors, mtD2 was detected in CD11b+ myeloid cells (**Fig**. **S1H**). Similarly, we also restricted mtD2 expression in myeloid cells by crossing PhAM^floxed^ with CD68^Cre^ (PhAM^+/+^CD68^Cre^) (**Fig S1I**). Tumor implantation resulted in mtD2 expression in CD11b^-^CD3^+^ lymphoid cells within the TME (**Fig S1J).** Altogether, the results demonstrate that mitochondria may be transferred from one immune cell type to others, a process orchestrated by cancer cells.

To demonstrate mitochondria redistribution by human cancer cells, we used mitotracker green dye (MTG) to label the mitochondria of T cells from a donor (APC labeled). Separate T cells from a different donor were differentially-labeled (CD3-PE). These two distinct T cell populations were co-cultured alone or in the presence of the melanoma line SK-MEL-28 directly or in transwell. Only in the direct presence of tumor cells did we observe mitochondria transfer between T cells (**Fig S1K**). Furthermore, we tracked mitochondria redistribution by cancer cells in human cancer patients, leveraging the computational mitochondria transfer inference program, MERCI^25^. Analysis of previously-validated human tumor scRNA-seq datasets revealed the redistribution of CD8^+^ T cell–derived mitochondria to multiple immune populations, including macrophages (**Fig. 1L**), dendritic cells, and B cells across multiple tumor types (**Fig S1L**). These findings establish mitochondrial redistribution as a conserved feature of both murine and human tumors.

### Mitochondria fusion in cancer cells prior to redistribution

The fate of mitochondria acquired from donor cells is uncertain; it is unclear if they persist as separate entities or integrate into the host cell’s mitochondrial network ^24,29^. To address this in cancer cells, we labeled *in situ* tumor mitochondria network with mitotracker red (MTR), and co-cultured them with CD45.2^mtD2+^ leukocytes. Confocal microscopy revealed that the vast majority (>80%) of exogenous mtD2 mitochondria fused seamlessly with *in situ* endogenous tumor mitochondria (**Fig. 2A–C**). We frequently observed concatenated hybrid structures containing both red and green signals, indicating physical integration of the two pools (**Fig 2B**).

**Figure 2.**
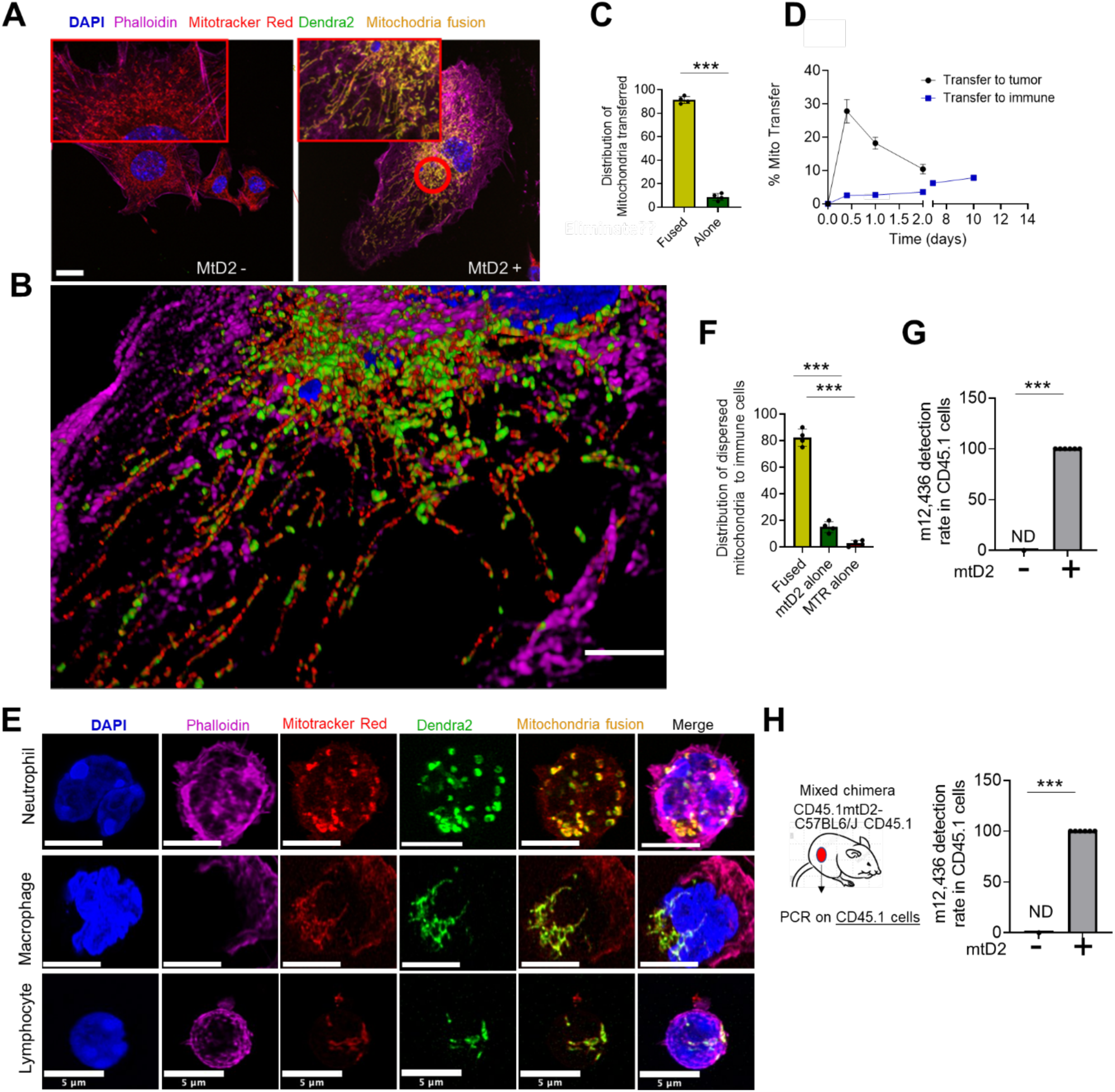
Mitochondria are fused prior to redistribution. **A**) Representative immunofluorescence staining tumor cells without or with exogenous mitochondria. **B**) 3-dimensional rendition of cancer cell with fused mtD2 (green) and endogenous mitochondria (red). **C**)Assessment of exogenous mtD2 mitochondria that are fused (attached to endogenous mitochondria) or alone within cancer cell. **D**) Rate of endogenous mitochondria transfer from cancer cells to immune cells (blue) and rate of mitochondria transfer from immune cells to cancer cells (black). **E**) Representative immunofluorescence images of CD45.1 neutrophils, macrophages and lymphocytes with fused exogenous mitochondria obtained from B16 cancer cells. **F**) Quantification of mitochondria dispersed from cancer cells to immune cells. **G**) Detection rate of m12,436 in CD45.1 co-cultured with cancer cells harboring exogenous mtD2. **H**) Detection rate of m12,436 in CD45.1 cells from CD45.1^mtD2-^/CD45.2^mtD2+^ mixed chimeric mice challenged with tumor. ***<0.001 by unpaired T test except f (ANOVA).

We next asked whether fusion was mechanistically linked to redistribution. Prior reports suggested that tumor-to-CD8 T cell mitochondria transfer is relatively inefficient compared to immune-to-tumor transfer ^16,25,30^. Consistent with this, we found that cancer cells exported mitochondria at lower frequency and slower kinetics than they imported them (**Fig. 2D**). After MTR+ B16: CD45.2^mtD2+^ leukocyte co-culture, we sorted mtD2+ cancer cells and co-cultured them with CD45.1^mtD2-^. Confocal microscopy revealed transfer of fused mitochondria to CD45.1 cells (**Fig 2E**). The vast majority (∼85 %) of mitochondria redistributed to CD45.1 cells were fused, mtD2+MTR+ hybrid mitochondria. About ∼12% of cells were only mtD2 positive, and rare CD45.1 cells (∼< 3%) were only MTR positive (**Fig 2F**). These findings indicate that cancer cells redistribute mitochondria not as isolated units, but predominantly as fused composites of endogenous and exogenous origin.

To further validate the dispersion of fused mitochondria by cancer cells, we employed a melanoma line engineered by base editing to carry an m.12436G>A mutation^31^. We performed triple co-culture with CD45.1^mtD2-^ and CD45.2^mtD2+^. We reasoned that if cancer dispersed fused mitochondria, m.12,436G>A mutation would be detected in CD45.1 immune cells that become mtD2+. Indeed, we detected the cancer cell-derived m.12,436G>A point mutation in CD45.1 cells, but only in the mtD2+ subset (**Fig 2G**). To demonstrate the dispersion of fused mitochondria *in vivo*, we implanted m.12436^80%^ tumors into mixed CD45.2^mtD2+^/CD45.1^mtD2-^ chimeric mice. After tumor explant, we gated on CD45.1 cells and could detect m.12436 mutation in the mtD2+ subset (**Fig 2H**), similar to our *in vitro* studies. Similarly, we subcutaneously implanted m.12,436G>A m tumors into CD45.2^NZB^/CD45.1^C57BL6/J^ mixed chimeric mice. CD45.1 cells were isolated, and we could confirm m.12436 mutation and NZB mtDNA in those cells (**Fig S2A**). These multiple lines of evidence indicate that cancer cells disperse fused mitochondria – a mixture of their own endogenous and exogenously-acquired mitochondria – to immune cells.

### Mechanisms of mitochondria redistribution

Among the various mechanisms enabling mitochondrial transfer to cancer cells, tunneling nanotubes (TNTs) play the dominant role, whereas extracellular vesicles account for a modest, though measurable, proportion of transfer events^32^ ^15,16,19^. Therefore, we sought to determine whether TNTs similarly mediate dispersion of fused mitochondria from cancer cells to immune cells. Indeed, super resolution imaging revealed actin-rich, cellular projections between cancer cells and immune cells, containing fused mitochondria (**Fig S2B**), consistent with TNTs. Less frequently, we also identified transfer of anuclear, mitochondria-laden vesicular detachments from cancer cells (**Fig S2C**), consistent with extracellular vesicles. To determine the relative contribution of these modalities to mitochondria redistribution from cancer, we performed initial tumor: CD45.2 leukocyte co-culture to allow mtD2 transfer to tumor cells. Then, after washing away immune cells, we introduced CD45.1 immune cells as we described previously, or under 0.4 µm filter to block contact, or in the presence of cytochalasin B to block TNTs, or GW4869 to block EV release. Inhibiting physical contact and TNTs substantially reduced mitochondria redistribution to immune cells by more than 85% By contrast, GW4869 only caused ∼15% reduction in mitochondria redistribution to immune cells (**Fig S2D**). These results indicate that TNT formation between cancer cells and immune cells facilitate the majority of mitochondria redistribution.

### Redistributed mitochondria reprogram acceptor immune cells

Next, we evaluated the functional consequence of mitochondria redistribution to recipient immune cells. Mitochondria are redistributed to many cell types (**Fig 3A**). We focused on neutrophils, macrophages, and CD4 T cells because they are the dominant immune infiltrates in tumors that physically interact with tumors and contribute to the immunosuppressive TME^33,34^. After mtD2+ tumor: CD45.1 immune co-culture, we distinguished recipient CD45.1 cells on the basis of mtD2 and compared their phenotype with mtD2- counterparts (**Fig 3B-D**). Neutrophils that acquired mitochondria (CD45.1+Ly6G+CD11b+mtD2+) showed increased pro-tumoral features, including elevated CD200R, PD-L1, and NETs formation (**Fig 3B**). Similarly, mtD2+ macrophages displayed higher levels of CD200R, PD-1, MerTK, and CD206 (**Fig. 3C**). In CD4 T cells, mitochondrial uptake promoted the expansion of FoxP3+CD25+ regulatory T cells (Tregs) (**Fig. 3D**). These Tregs exhibited heightened immunosuppressive activity, with increased PD-1, CD69, and FoxP3 expression (**Fig. 3E**), enhanced IL-10 production (**Fig. 3F**), and stronger suppression of target T cell proliferation (**Fig. 3G**). Together, these data indicate that mitochondria redistributed from tumor cells reinforce immunosuppressive programs across multiple immune cell types.

**Figure 3.**
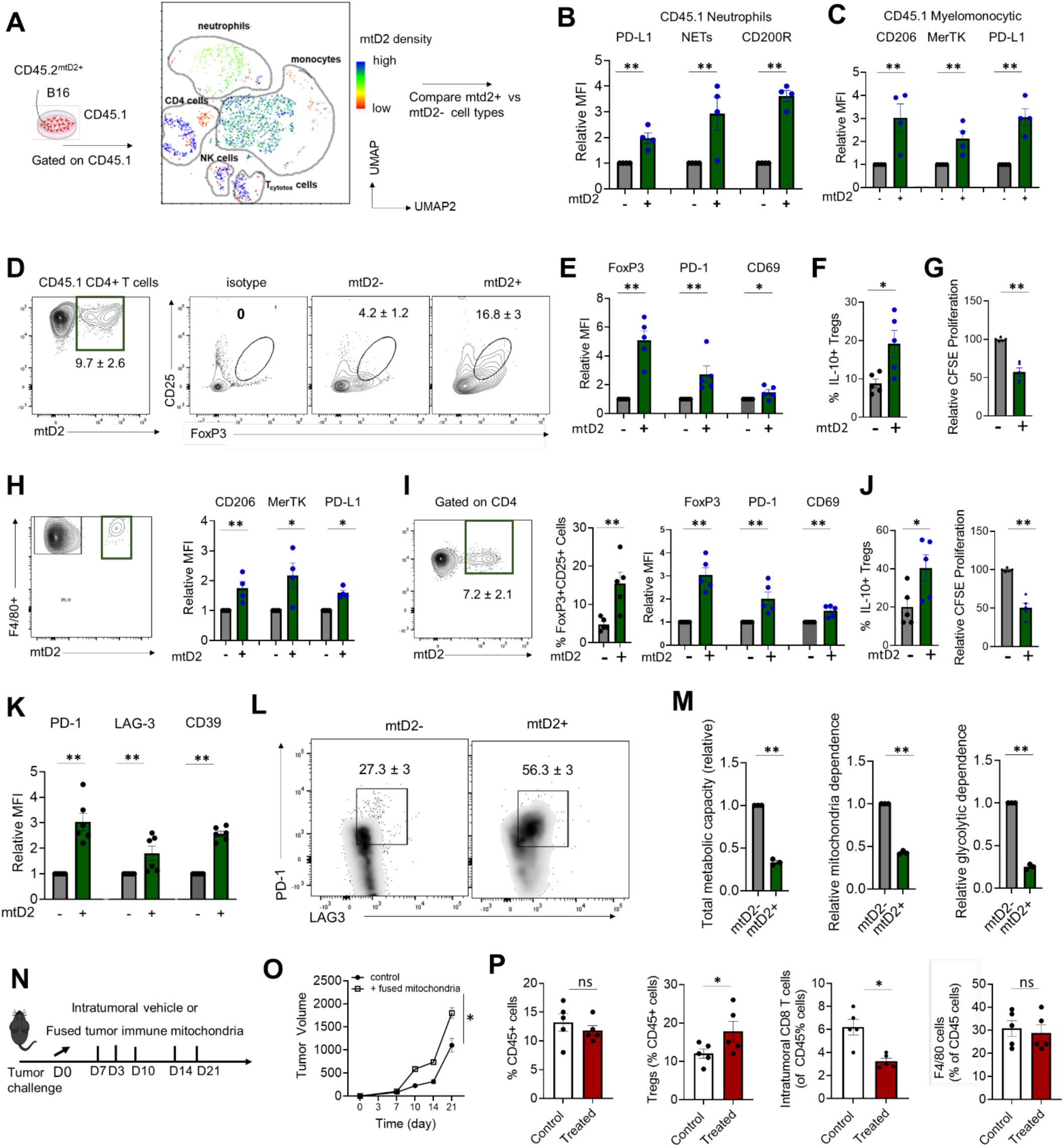
Dispersed mitochondria reprogram acceptor immune cells. **A)** Schematic for B16: CD45.2^mtD2^ co-culture followed by redistribution of mitochondria to CD45.1 cells. **B**) Relative expression of PD-L1, NETs and CD200R in CD45.1 neutrophils that accept mtD2 versus neutrophils that do not. **C**) Relative expression of CD206, MerTK, PD-1 in macrophages that accept exogenous mtD2 versus mtD2- macrophages. **D**) Expression of Foxp3, CD25+ Tregs in CD4 cells that acquire exogenous mtD2 mitochondria. **E**) Relative expression of FoxP3, PD-1 and CD69 in Tregs that have accepted exogenous mitochondria (mtD2+) or not (mtD2-). **F**) IL-10 secretion and suppressive capacity. **G**) of mtD2+ and mtD2-Tregs. **H-I**) Phenotype of CD45.1 macrophages and Tregs in vivo from CD45.2^mtD2^ and CD45.1^mtD2-^ mixed chimeras inoculated with tumors. Relative expression of CD206, MerTK and PD-L1 by mtD2+ and mtD2- F4/80 macrophages (CD45.1+ cells). **J**) Relative IL-10 production and proliferative suppression of target cells in mtD2+ and mtD2- Tregs in vivo. **K-M**) Profile of CD8+ T cells that accept mitochondria from tumors versus those that did not. K shows expression of exhaustion markers on mtD2+ CD8 T cells relative to control. **L)** shows percentage of CD8+ T cells that express PD-1 and LAG3. **M)** shows SCENITH metabolic profiling of metabolic capacity, mitochondria and glycolytic capacities. **N)** Schematic for tumor inoculation and assessment of tumor growth after intratumoral administration of fused mitochondria. **O**) B16 tumor growth in mice administered with vehicle or fused mitochondria (n=8 per group). **P**) Frequencies of total immune cells, Tregs, CD8 T cells, and macrophages in control versus mitochondria administered tumors. p<0.05, **p<0.01, ***p<0.005. NS indicates not significant. Statistical analyses were by Student’s test; O by two-way ANOVA followed by Bonferroni correction.

To further examine the direct role of exogenous fused mitochondria as a causal driver of immunosuppression, we prepared tumor: leukocyte co-cultures and harvested their mitochondria. Then, using mitoception protocol^35^, we directly introduced the fused cargo into different immune cells. Macrophages differentiated in the presence of exogenous fused mitochondria increased expression of CD206, MerTK, similar to our observation from direct dispersion of mitochondria from cancer cells (**Fig S3A**). Differentiation of CD4T cells in the presence of fused mitochondria also enhanced FoxP3, CD25, and PD-1 expression (**Fig S3B**).

To characterize the phenotype of immune cells that accept fused mitochondria *in vivo*, we turned to the mixed CD45.2^mtD2^/CD45.1^mtD2-^ mice challenged with B16 tumor. We gated on CD45.1 immune cells, and subgated macrophages and CD4 T cells on the basis of mtD2 status (**Fig 3H** and **Fig S3C** for gating strategy). F4/80+ CD45.1 cells that acquire exogenous mitochondria upregulated PD-L1, MerTK and CD206 (**Fig 3H**). CD4+ CD45.1 T cells that acquire exogenous mitochondria upregulated PD-1, FoxP3 and CD25 (**Fig 3I**). They produced more intracellular IL-10 and suppressed the proliferation of target cells with greater potency (**Fig 3J**). Thus, transfer of mitochondria also enhances immunosuppressive features *in vivo*.

Beyond immunosuppressive cells, CD8+ T cells are also frequent recipients of redistributed mitochondria (**Fig 3A**). We found that CD8+ T cell recipients of tumor-redistributed mitochondria showed evidence of exhaustion, including upregulation of PD-1, LAG3, and CD39 (**Fig 3K**) as well as a higher frequency of PD-1^hi^LAG^hi^ cells (**Fig 3L**). Moreover, using SCENITH metabolic analysis^36^, we found that these CD8+ T cells demonstrated marked decrease in total metabolic capacity, characterized by decreased mitochondria and glycolytic capacities (**Fig 3M**). Thus, while mitochondria redistribution enhances suppressive cells, it leads to functional and metabolic exhaustion of CD8+ T cells.

To determine the impact of mitochondria dispersion on cancer outcomes, we subcutaneously challenged mice with B16. When tumors were established, we harvested fused mitochondria derived from tumor: immune co-cultured and directly injected them into the tumor bed (**Fig 3N**). Compared to tumors treated with vehicle, those injected with fused mitochondria showed a more rapid growth rate (**Fig 3O**). Direct injection of mitochondria into the tumor bed did not change the frequency of immune cell infiltration into the tumors. However, this expanded the frequencies of Tregs, decreased the frequency CD8+ T cells, but had no impact on the frequency of infiltrating macrophages (**Fig 3P**). Conversely, we treated cohorts of B16 tumor-bearing mice with inhibitors of mitochondria transfer using a combination of EV inhibitors (GW4869) and TNT inhibitors (L-778123)^16^ (**Fig 3SD**). Such therapy reduced tumor growth, limited the expansion of Tregs and increased the frequency of CD8+ T cells (**Fig S3E).** Therefore, mitochondria redistribution can augment tumor growth, whereas inhibition of mitochondria transfer can restrain tumor growth.

### Mitochondria fusion optimizes metabolic fitness of cancer cells

Multiple recent reports indicate that mitochondria transfer can expand regulatory T cells ^37–39^. In those studies, the transferred mitochondria were not fused with tumor mitochondria. Therefore, the dispersion of fused mitochondria by cancer cells may not be required to induce the expansion of regulatory cells. On that basis, we speculated that cancer cells undergo mitochondria fusion for metabolic gains. To explore this possibility, we performed RNAseq of cancer cells with fused mitochondria. Compared to their respective controls, B16 or E0771 cancer cells with fused mitochondria had markedly increased mitochondrial macromolecular biosynthetic capacity (**Fig 4A).** Membrane potential, measured by TMRM assay, was increased in mtD2+ cancer cells **(Fig 4B).** SCENITH metabolic analysis revealed that mtD2+ cancer cells substantially increased their total metabolic capacity, largely driven by enhanced mitochondria dependence (**Fig 4C**). These results suggested that exogenous mitochondria augment the mitochondrial macromolecular biosynthetic fitness of cancer cells.

**Figure 4.**
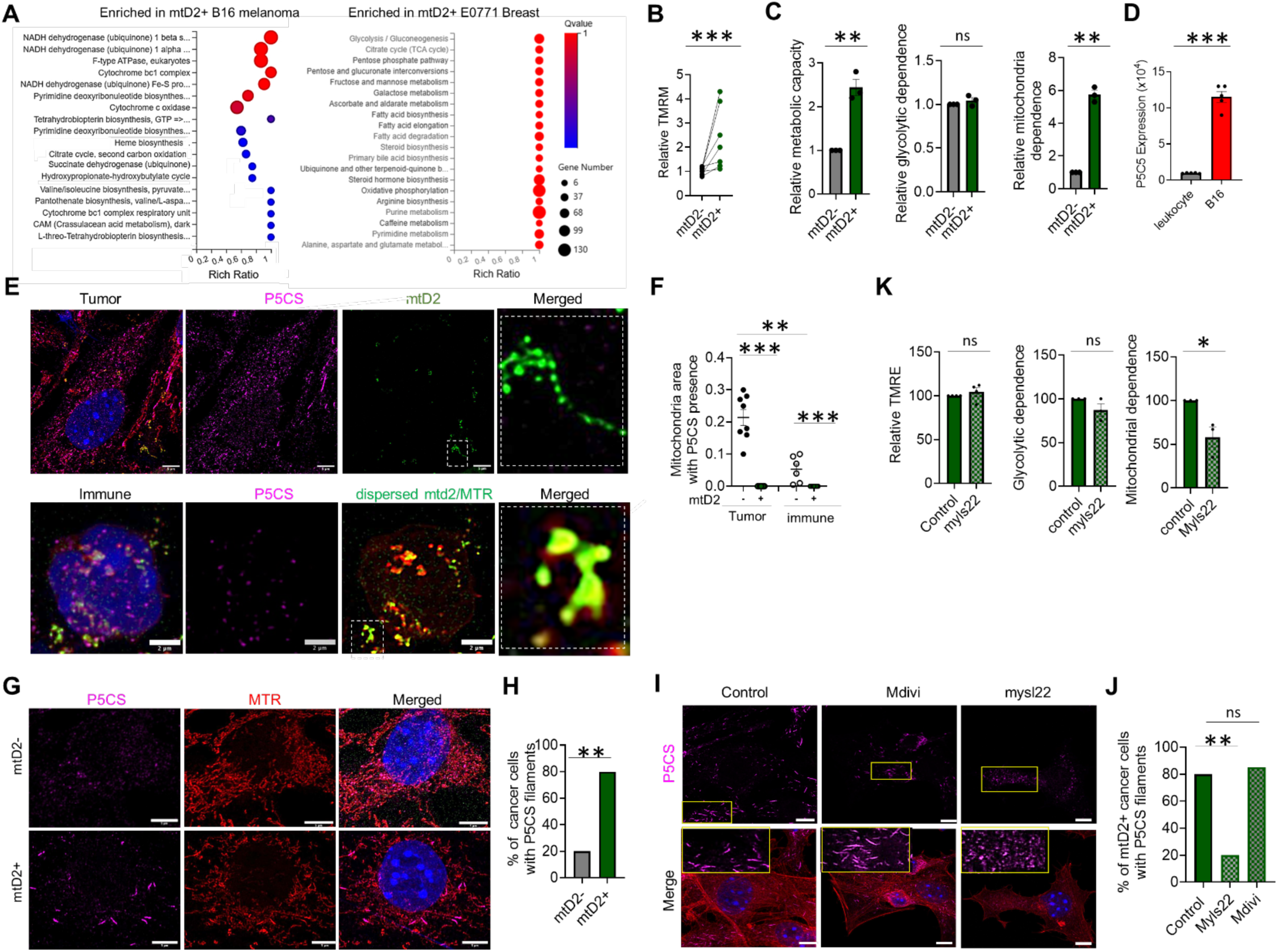
Mitochondria dispersion improves metabolic fitness of cancer cells. **A**) Metabolic pathway enrichment in tumor cells with exogenous mitochondria. n= 3 independent pairs of mtD2+ and mtD2- cancer cells from the same experiment. **B**) Membrane potential mtD2+ vs mtD2- cancer cells. n=5 matched cohorts. **C**) Metabolic profiling of mtD2+ vs mtD2- B16 cancer cells by SCENITH. **D**) P5CS expression levels in leukocytes and melanoma cells. **E)** Representative immunofluorescence of P5CS staining of cancer cells that have acquired mtD2 vs mtD2- cancer cells. P5CS does not colocalize with mtD2. Bottom panel: Immunostaining of P5CS expression in accepting immune cells. P5CS does not colocalize with transferred, fused mitochondria. **F**) Pearson correlation coefficient analysis for mitochondria areas overlapping with P5CS in tumor cells with or without mtD2, and in immune cells that have received redistributed mtD2 or not. **G**) Immunofluorescence images of P5CS conformation in mtD2+ vs mtD2- cancer cell. **H**) Quantification of the frequency of mtD2+ and mtD2- cancer cells with filaments P5CS conformation. **I**) Representative images of P5CS architecture in cancer cells with exogenous mitochondria (mtD2) in the presence of myls22 or mdivi-1. **J**) Quantification of P5CS filament in mtD2+ cancer cells treated with fusion or fission inhibitors versus control. **K**) Effect of blocking mitochondria fusion on metabolic capacity of mtD2+ cancer cells. *p<0.05, **p<0.01, ***p<0.005. NS indicates not significant. Statistics were performed using paired t test (B), unpaired t test (C) and ANOVA (F). bar indicates 5 µm.

Under various bioenergetic demands, mitochondria can segregate into two conformations to enable them execute reductive synthesis and oxidative phosphorylation simultaneously. This is mediated by compartmentalization of mitochondria into two subpopulations: a canonical cristae-rich architecture, which enables complex V docking for ATP production; and a fibrillar architecture, which favors macromolecule biosynthesis. The latter conformation is associated with the sequestration of pyrroline-5-carboxylate synthase (P5CS) filaments, which drives reductive biosynthesis^22^. Therefore, we hypothesized that transferred mitochondria trigger P5CS filamentous structural changes in recipient cancer cells.

We first tested whether cancer cells utilize mitochondria transfer to import P5CS-rich mitochondria from host immune cells. Measurement of P5CS levels in leukocytes compared with tumor cells showed more than a log-fold less P5CS expression intensity in leukocytes than in cancer cells (**Fig 4D**). Although tumor cells express abundant P5CS, we failed to colocalize P5CS with any of the transferred mitochondria in mtD2+ cancer cells (**Fig 4E**, and **Fig S4A**). Conversely, when mtD2+ cancer cells redistribute mitochondria to CD45.1 acceptor cells, P5CS is not co-transferred (**Fig 4E,F**). These results suggest that cancer cells do not import P5CS-rich mitochondria and most likely exclude P5CS from the transferred mitochondrial pool. Indeed, net P5CS increase in cancer cells after they disperse fused mitochondria is minimal (**Fig S4B**) and P5CS levels do not change in immune cells that accept fused mitochondria (**Fig S4C**). Strikingly, however, we found that exogenous mitochondria triggered fibrillar conformation of P5CS in recipient cancer cells (**Fig 4G,H**), and many cancer cells with exogenous mitochondria showed fibrillar P5CS re-organization in clusters (**Fig S4D**).

P5CS architectural changes are commonly driven by cellular stress responses^22,40,41^. Therefore, we asked whether the cellular stress associated with mitochondria fusion directly contributes to the filamentous architecture change in cancer cells after acquiring exogenous mitochondria. To that end, we performed B16: CD5.2^mtD2+^ co-cultures as we have described or in the presence of Myls22, a small molecule inhibitor of OpA1^42,43^, which governs mitochondria inner membrane fusion^44^. As an additional control for the potential mitochondrial dynamic changes that could be induced by Myls22, we treated separate tumor: immune co-culture cohorts with Mdivi-1, which promotes mitochondria fusion through Drp1.^45,46^ Disrupting endogenous and exogenous mitochondria fusion in cancer cells prevented the filamentous morphological changes in P5CS, whereas treatment with Mdivi-1 did not (**Fig 4I,J**). Importantly, this disruption did not impact membrane potential but reduced the mitochondria-specific metabolic gains in mtD2+ cancer cells (**Fig 4K)**. These results indicate that the fusion of exogenous mitochondria with endogenous network promotes P5CS architectural organization, contributing to the enhanced metabolic fitness in cancer cells that hijack and redistribute mitochondria.

Finally, we asked whether fusion of exogenous mitochondria can enable tumors to utilize dysfunctional, host-derived mitochondria. We created mitochondria reporter mouse on the background of mice lacking Ndufs4 (Nfuds4KO^mtD2^). Ndufs4 encodes mitochondria complex I and is essential for cellular energy production. Interestingly, the rate of mitochondria transfer from Nduf4^KO^ and Nduf4^WT^ leukocytes were similar (**FigS4E**). The mitochondria transfer from Nduf4^KO^ leukocytes still improved membrane potential and enhanced the bioenergetics of recipient cancer cells to the same extent as transfer from Nduf4^WT^ leukocytes (**FigS4F, G**). Thus, mitochondria fusion enables cancer cells to repurpose suboptimal mitochondria to promote tumor metabolism.

## DISCUSSION

Vast bioenergetic and biosynthetic resources are required to sustain tumor growth. Mitochondrial metabolism, in particular, fuels cancer cell proliferation and is also required by immunosuppressive cells to direct antitumor immunity^7,9,13,47^. Yet, the TME is hostile to mitochondrial metabolism. The highly mobile nature of mitochondria^48^, and the principles of maximum parsimony raised the prospects that a coordinated mechanism might exist to sustain the mitochondrial metabolism of cancer cells and their immunoregulatory partners. Here we uncover such a mechanism: cancer cells import exogenous mitochondria, fuse them with their own networks, and redistribute the fused payload to neighboring immune cells to fuel the immunosuppressive TME. Mitochondria redistribution has divergent effect on recipient immunosuppressive and cytotoxic CD8 T cells, as it boosts the expansion of the former and exhausts the latter. One possible explanation is that the transferred mitochondria accumulate reactive oxygen species^24^. Whereas regulatory cells possess abundant bespoke antioxidant systems and are redox-resilient, cytotoxic effector CD8 T cells are relatively redox-vulnerable^49–53^. The fusion of exogenous and endogenous mitochondria also reconfigures tumor cell metabolism through P5CS conformational remodeling, thereby allowing cancer cells to couple ecosystem regulation with tumor-intrinsic metabolic gain.

The ability of relatively small amounts of transferred mitochondria to dramatically boost the metabolism of recipient cells has been met with much skepticism^54,55^. Our research shows that, at least in cancer, this can be achieved by P5CS architectural changes in the endogenous mitochondrial network. Whether such mechanisms contribute to the metabolic boost in other cells under various clinical and pathologic settings following transfer remains to be elucidated.

The fact that cancer cells acquire and redistribute mitochondria in a fused form has several implications. This axis provides extraordinary metabolic flexibility. As we showed, cancer cells can even incorporate suboptimal mitochondria and still extract metabolic benefit. The maneuver could also enable tumors to selectively offload dysfunctional mitochondria to shape their surroundings. An example of the latter has recently been reported, wherein cancer cells transferred *in situ* mtDNA mutation to recipient CD8 T cells, resulting in impaired immune function^30^. The incorporation of fresh, host-derived mitochondria along with the expulsion of *in situ* mitochondria may provide mechanistic basis for how cancer cells, replete with mtDNA mutations^31,56^, avoid catastrophic collapse of mitochondrial function: they can exploit beneficial mutations while replenishing their network with host-derived mitochondria.

Mitochondria transfer is frequently associated with dampening immune response and promoting tolerance in various settings. For instance, it improves inflammation resolution in the lung following pulmonary injury^57,58^. In diabetic renal injury, it restricts macrophage inflammation^59^. Mitochondria transfer from mesenchymal cells also induces potent Tregs and suppressed Th1 proliferation ^37–39^. Consistent with those reports, cancer cell redistribution of fused mitochondria instructed immune cells in the TME towards immunosuppressive fates. The fundamental underpinning of mitochondria’s effect on immunity remains unclear. We speculate that sharing mtDNA may blur the genetic boundary between tumor and host, increasing camouflage, tolerance, and immune evasion. This perspective of organelle sharing evokes mitochondria symbiosis, the foundational, primordial act of cooperation that beget eukaryotic life, when proto mitochondria integrated with Asgard archaea ^60,61^. Thus, we propose that redistributed mitochondria from cancer do not only provide metabolic substrate for the expansion of regulatory cells, but that sharing hybrid mitochondria may regulate the TME by communicating cellular co-existence within the tumor ecosystem.

In summary, we define mitochondrial redistribution as a mechanism by which tumors simultaneously reinforce their own biosynthetic capacity and construct an immunosuppressive niche. Rather than passive metabolic scavengers, cancer cells are active distributors of organelles, coordinating growth with ecosystem control. Mitochondria redistribution may represent a central organizing principle of tumor biology, opening up new therapeutic levers distinct from current metabolic or checkpoint-based strategies. Given the deleterious impact on CD8 T cells, it may be clinically important to determine whether cancer-redistributed, fused mitochondria contribute to the exhaustion of CAR-T cells upon infiltrating the TME.

### Limitations

We probed numerous publicly available scRNAseq datasets of cancer patients from the CancerSCEM database^62^. Unfortunately, many datasets were not candidates for computational inference of mitochondria redistribution using MERCI. This is largely attributed to insufficient coverage of mtDNA reads that failed to meet the rigorous threshold necessary for accurate prediction of mitochondria transfer from CD8^+^ T cells to other cells besides cancer cells (see Methods for details). Moreover, MERCI is only currently trained to predict mitochondria transfer from CD8^+^ T cells. As such, this limits our ability to extensively infer mitochondria redistribution in real world cases of human cancers. The emergence of mtscATACseq will enable more accurate prediction of mitochondria redistribution in solid cancer to understand the full impact of this mechanism in human disease.

## MATERIALS AND METHODS

### Mice

Wildtype C57BL6/J CD45.1 (strain #000664), C57BL6/J CD45.2 (strain # 002014), NZB (strain #:000684), mtDendra2^Flox/Flox^ (PhAM Flox; also referred to as mtD2, strain #18385), PhAM^excised^ (global mtD2 mitochondria reporter, strain #:018397), CD4^Cre^ (strain # 022071), CD68^CreERT2^ (strain #:038175), CD8Cre (strain # 008766), Nduf4-deficient mice (strain #:027058) were purchased from The Jackson Laboratory (Maine, USA) and bred in-house for experimental use. The global mitochondria reporter mice are on the CD45.2 background (CD45.2^mtD2^). To generate CD45.1^mtD2^, mtD2 mice were crossed for more than 10 generations with CD45.1 to eliminate the CD45,2 allele. These mice were a generous gift from the Brestoff Laboratory (Washington University, St Louis). All mice had *ad libitum access* to food and water and were maintained in a specific-pathogen-free facility with a 12h:12h light: dark cycle (lights on from 0600 to 1800). Experiments were performed in male and female mice between 8 and 12 weeks. Animals were randomly assigned to groups per experiment, ensuring similar distribution of age and gender. No specific blinding method was used. Data represent at least 2 independent experiments that were pooled for analysis. All experiments were executed under the guidelines of the Institutional Animal Care and Use Committee (IACUC # 34210).

### Bone Marrow Chimera

Briefly, recipient mice received two doses of 5.5 Gy irradiation 3 hours apart. Thereafter, femurs of donor mice (CD45.1^mtD2-^ and CD45.2^mtD2+^) were flushed. Red blood cells were lysed and equal numbers of cells from each donor pool were mixed (total 10 x10^6^ cells) and intravenously administered to irradiated recipient mice. Blood, spleen, and marrow were sampled from mixed chimeric mice to confirm successful engraftment prior to experimental use.

### Parabiosis

Parabiosis was performed as previously described^63^. Parabiosis was performed using 7-week-old female mice, with each pair consisting of a wildtype CD45.1^mtD2-^ and CD45.2^mtD2+^ mice. Mice were anesthetized with 2–3% isoflurane and placed on a sterile surgical field. The corresponding lateral aspects (left for one mouse, right for the other) were shaved, cleaned with 70% ethanol and povidone-iodine, and incised from elbow to knee to expose the underlying musculature. The skin was gently loosened from the subcutaneous tissue along the flank. The olecranon and knee joints of each mouse were ligated together using nonabsorbable 5-0 silk sutures to ensure musculoskeletal alignment. The dorsal and ventral skin were then sutured together using 6-0 nylon monofilament in a continuous or interrupted fashion. Following surgery, mice were maintained on a heating pad and monitored continuously until fully ambulatory. Analgesia was administered with sustained-release buprenorphine (1.0 mg/kg subcutaneously) and meloxicam (5 mg/kg) as needed. Pairs were housed in large cages with accessible food and water gel. Circulatory cross-engraftment was confirmed by flow cytometry for CD45.1/CD45.2 chimerism.

### In vivo animal studies (drug and mitochondria treatment)

B16 melanoma cancer cells (1 × 10^6^ cells) were subcutaneously injected in the flanks of syngeneic mice. The drug therapy was started on day 3 post tumor implantation. Each animal was intraperitoneally injected every alternate day with the vehicle (for the control group), αPD1 (10 mg kg^−1^) alone, combined L-778123 (80 mg kg^−1^) and GW4876 (3 mg kg^−1^) or a combination of all three drugs αPD1 (10 mg kg^−1^, L-778123 (80 mg kg^−1^), and GW4876 (3 mg kg^−1^). The tumors were measured at indicated times using a Vernier caliper, and the tumor volume (*V*_t_) was calculated as per the following formula: *L* × *B*^2^/2, where *L* is the longest dimension and *B* is the shortest dimension. The total body weight was routinely measured to assess any gross toxicity. All the tumor tissues were harvested for further studies. The maximum permitted tumor volume (2 cm^3^) was not exceeded in any study.

For adoptive mitochondria transfer, mitochondria were isolated from cancer cell and immune cell co-culture using the mitochondria isolation kit (Thermo Fisher Scientific, cat# PI89874). We typically prepared mitochondria from a co-culture of 1x10^7^ and 1x10^7^ blood leukocytes per mouse. Purified mitochondria were injected directly into tumor beginning day 3.

### Cancer cells, culture and treatments

Tumor cells (B16F10, E0771, MC38, SK-MEL-28 were originally obtained from ATCC. Cells were retrovirally transduced to overexpress Tdtomato. Cells were cultured according to manufacturer’s conditions, typically consisting of RPMI 1640 or DMEM, 10% heat inactivated fetal bovine serum (FBS) and 1% penicillin and streptomycin. Cell lines are authenticated through STR profiling, and all lines were subjected to mycoplasma testing every 6 months using the Universal Mycoplasma Detection Kit.

For in vitro co-cultures, tumor cells were seeded in 24-well plates with immune cells, typically using 1 x10^5^ tumor cells in1:1 ration unless otherwise specified. After 16 h or indicated time, cells were washed and evaluated for mitochondria transfer to tumor cells. For mitochondria dispersion, cancer cells were sorted from immune cells (CD45.2) and re-cultured with immune cells harvested from CD45.1 mice.

For in vivo experiments, cells were implanted subcutaneously on the flanks of recipient mice at 1 x10^6^ in 50 ul PBS. Between 14-21 days, tumors were explanted, digested in collagenase IV, and passed through 70 µm strainer to obtain single cell suspensions. Flow cytometry was then used to identify mtD2 expression by live tumor cells (Tdtomato^+^CD45^-^CD31^-^DAPI^-^).

### Mitoception

Mitoception was performed as previously described^35^. Briefly, mitochondria from tumor: immune co-culture were harvested and prepared in a 500 µl suspension in DMEM/FCS 5%. The suspension was added slowly to the plated recipient cells, close to the bottom of the well, throughout the culture surface. The culture plates were then centrifuged at 1,500 g for 15 min at 4°C. They were then placed in a 37°C cell incubator prior to a second centrifugation in the same conditions, two hours later, and then returned to 37°C cell culture.

### Confocal Microscopy

Cancer cells (1 x10^4^) were seeded on # 1.5 round coverslip prior to the addition isolated mtD2 immune cells in a 1:1 ratio. The cells were co-cultured on the coverslip for indicated times at 37 °C in 5% CO2 incubator. The cells were then fixed with 4% paraformaldehyde and kept at 4°C until staining. The samples were stained with 300 nM DAPI, Alexa Fluor 647 phalloidin (1:500), and anti-mouse CD45-PE (1:500) for 30 minutes, at room temperature, followed by two times washing with PBS. The images were acquired with a Zeiss LSM900 confocal microscope with Plan-Apochromat 63X/1.4 oil objective.

Co-cultures of cancer and immune cells were imaged using a Leica Stellaris 5 White Light Laser (WLL) confocal microscope equipped with a motorized stage and a Leica DMi8 inverted microscope base. The system included a 405 nm solid-state laser and a tunable White Light Laser (WLL) providing excitation wavelengths from 485 nm to 790 nm. Fluorescence emission was detected by highly sensitive Power HyD S detectors, with spectral detection ranges optimized for each fluorophore to minimize crosstalk. All images were acquired with a 63x NA 1.4 oil immersion objective.

Super-resolution imaging was achieved using the integrated LIGHTNING adaptive deconvolution module within the Leica LAS X software. This algorithm adaptively processes the image data by determining optimal deconvolution parameters for each voxel based on local image properties, thereby increasing image resolution and contrast beyond the diffraction limit. The deconvolution process was applied automatically during or immediately after acquisition to generate the super-resolution images presented throughout the manuscript. Original raw confocal images are also available. All image processing, including brightness, contrast adjustments, and 3D rendering, was performed using Leica LAS X software and ImageJ/Fiji (National Institutes of Health, USA).

Antibodies were used at a dilution of 1:1000 for immunofluorescence unless otherwise indicated. For multiplexed immunostaining, we used highly cross-adsorbed secondary antibodies to avoid cross-species contamination. The following antibodies were used: Phalloidin-iFluor™ 647 Conjugate (Cayman Chemicals, Cat # 20555), anti-rabbit IgG alexa fluor 647 (Highly cross-adsorbed, Invitrogen, A-31573; 1:500 for immunofluorescence), Mitotracker Red (ThermoFisher), P5CS (Proteintech, 68184-1-Ig, 1:500 for immunofluorescence).

### Flow Cytometry

For flow cytometric analysis of ex vivo tumors, Tumors were digested with collagenase IV and Dnase I and mashed through a 70 μm cell strainer (Falcon). After red blood cell lysis, cells were then stained with antibody cocktail in FACS buffer (PBS with 0.5% bovine serum albumin) at 4 degrees for 30 minutes. Thereafter, staining was quenched with washes with washing with FACs buffer and immediately analyzed on a flow cytometer. Antibody cocktails included DAPI or 7-AAD to exclude dead cells. Samples were analyzed with FlowJo software 10.8.2 (BD Biosciences). The following antibodies (BioLegend) were used: anti CD45 (clone 30-F11), anti-Ly6G (clone 1a8), anti CD11b (clone M1/70), anti-CD69 (clone H1.2F3) anti-CD3 (clone 145-2C11), anti-CD8a (clone 53-6.7), anti-PD-L1 (10F.9G2), anti-PD-1 (RMPI-30), were from Biolegend; LSR II or BDFortessa and BD FACSymphony (BD Biosciences) were used for flow cytometry acquisition and a FACSAria Fusion (BD Biosciences) or BD Influx (BD Biosciences) for cell sorting.

### SCENITH

Single cell metabolic analyses of cells were performed using SCENITH^36,64^ as previously described. In brief, after mitochondria transfer to cancer cells, or after mitochondria redistribution by cancer cells, each sample condition was treated with 10 µL of either wash media, 2-Deoxy-D-Glucose (2-DG, final concentration 100 mM), Oligomycin (Oligo, final concentration 1 μM), or a combination of 2-DG and Oligo. After 15 mins incubation at 37°, all conditions were treated with 10.5 µg/mL of puromycin, followed by a 40 mins, 37°C incubation. Cells were washed in 200 µL cold PBS, centrifuged 500 x g for 3 mins at 4°C and washed with FACS buffer. Zombie NIR viability dye, Fcblock, and surface staining was performed, followed by fixation, permeabilization and intracellular staining of puromycin using anti-puromycin (clone 12D10, Sigma-Aldrich, cat# MABE343-AF488). The samples were incubated 1 hr on ice, in the dark. Samples were washed with PermWash buffer (from the CytoFix/CytoPerm kit) and resuspended in 200 µL FACS buffer and subjected to flow cytometric analysis on a Cytek Aurora spectral flow cytometer configured with 4 lasers (violet, blue, yellow/ green, and red lasers), with 100 μL acquired to enable cell count enumeration.

### Chemicals

Tetramethylrhodamine (TMRE, Invitrogen, T669), Mitotracker deep red (Thermo, M22426), Myls22 (TargetMol, T9127); Mito-Tempol (Medchem express), GW4869 (Selleck Chemicals, S7609), Y27632 (Selleck Chemicals, S1049), cytochalasin B (Medchem Express, HY-16928).

### RNA seq

After tumor-immune co-cultures, cells were resuspended in FACS buffer (PBS in 0.5% BSA) and stained with DAPI and CD45. Live tumor cells (TdTomato+CD45-DAPI-) were sorted into mtD2+ and mtD2- subpopulations. Total mRNA was isolated using Qiagen RNEasy Plus isolation kit. Bulk RNAseq was performed using the NovaSeq platform. The raw RNA sequencing data was mapped to mouse reference genome hg38 using the STAR aligner, and genes annotated in Gencode v3664 was quantified using featurecounts in the subread package. The differential gene expression analysis was conducted in the DESeq2 package. Gene set enrichment analysis was performed with Gene Set Enrichment Analysis. Significantly different genes were identified by DESeq2 using Wald test. Gene annotation enrichment analysis was performed using KEGG pathways and GO terms (biological process, cellular component, and molecular function). Functional annotation clustering was performed and terms with p < 0.05 (Benjamini corrected) are shown.

### Mitochondria DNA PCR

After co-culture, mtD2- and mtD2- cells were FACS sorted. DNA isolation (Qiagen) following manufacturer’s instructions, qPCR reactions were designed with 100 ng of DNA and PowerUP SYBR Green Master Mix reagent (Thermo, A25742). Primers ARMS22 (5′-TTATCCACGCTTCCGTTACGTC-3′) and MT20 (5′-TGGCACTCCCGCTGTAAAAA- 3′) were used to amplify NZB mtDNA as previously described ^65^. NZB mtDNA (Ct) and the nDNA gene β-actin (Ct) were measured in samples obtained from tumors implanted into C57/NZB or WT/MtD2 chimeras or tumor cells treated with leukocytes isolated from either NZB or C57BL6/J mice. The NZB mtDNA signal was normalized to β-actin. Percentage of NZB mtDNA to the total mtDNA was calculated as NZB mtDNA/ (NZBmtDNA + C57mtDNA).

Bio-Rad QX200 Droplet Digital PCR System was used to quantify m34126 mtDNA in immune cells. Primers for m34126 are available as previously reported ^31^.

### MERCI

Computational inference of MERCI was performed as previously described^25^. To validate mitochondria redistribution in human samples, we first analyze scRNAseq datasets and filter samples with two minimal criteria: 1). Each sample contained at least 100 CD8+ T cells and 100 cancer cells (benchmarked for internal control), and 100 of any target cell of interest (example: macrophages). 2) the fraction of transferred mitochondria in cancer cells and the target cell of interest must be significantly associated with the gene expression variations on UMAP (|ρ| > 0.2, p < 0.001). DNA and RNA scores were estimated using mtDNA mutations and gene expression profiles as previously described^25^. Consistently, RNA and DNA scores with Rcm >1 cutoff was used as a criterion to select samples with mitochondria redistribution. We validate that receivers are sufficiently included in the input mixed cells. Samples that do not meet these criteria are excluded from downstream analysis. We then employ MERCI leave-one-out (LOO) pipeline to estimate the fraction of transferred mitochondria in target cells.

### Statistics

Data are reported as mean ± standard error of the mean. Statistical analyses were performed in Prism v10 or v11 (Graphpad, La Jolla, CA) unless otherwise specified. Paired or unpaired Student’s *t*-tests were used for two-group comparisons, with Welch’s correction applied when the standard deviations between groups differed. One-way analysis of variance (ANOVA) with Tukey or Fisher’s LSD post-hoc testing was used for three-or-more group comparisons, and two-way ANOVA with Bonferroni post-hoc. Statistical significance was set at p<0.05. Standard flow cytometry data were acquired and analyzed using a BD FACSymphony A3 flow cytometer and FlowJo software (BD Biosciences). Cell populations were sorted to >95% purity using BD FACSAria II (BD Biosciences) or Bigfoot Spectral Cell Sorter (Thermo Fisher Scientific).

## AUTHOR CONTRIBUTIONS

Conceptualization: DOD, AT

Design & Investigation: AT, ATW, VS, KB, YT, CCR, VS, CK, MM, LW, VRR, PG, TR, JDA, BL, EE, BL, DO-D

Formal Analysis: AZ, ATW, VS, YT, BL, DO-D

Writing – Original Draft- DO-D, EE, AT

Writing – Review & Editing - All

Supervision: DO-D, BL, JDA, EE

Funding Acquisition: DO-D

Funding: This project was supported by NIH R01ES034235 (DO-D), 5U54CA274511 (EE), 5P01CA244114 (EE), NIH R01CA258524 (BL).

Conflict: None

